# Synergistic survival-related effects of larval exposure to an aquatic pollutant and food stress get stronger during and especially after metamorphosis and shape fitness of terrestrial adults

**DOI:** 10.1101/2023.02.24.529881

**Authors:** Sarah Jorissen, Lizanne Janssens, Julie Verheyen, Robby Stoks

## Abstract

To improve the ecological risk assessment of aquatic pollutants it is needed to study their effects not only in the aquatic larval stage, but also in the terrestrial adult stage of the many animals with a complex life cycle. This remains understudied, especially with regard to interactive effects between aquatic pollutants and natural abiotic stressors. We studied effects of exposure to the pesticide DNP (2,4-Dinitrophenol) and how these were modulated by limited food availability in the aquatic larvae, and the possible delayed effects in the terrestrial adults of the damselfly *Lestes viridis*. Our results revealed that DNP and low food each had large negative effects on the life history, behaviour and to a lesser extent on the physiology of not only the larvae, but also the adults. Food limitation magnified the negative effects of DNP as seen by a strong decline in larval survival, metamorphosis success and adult lifespan. Notably, the synergism between the aquatic pollutant and food limitation for survival-related traits was stronger in the non-exposed adults than in the exposed larvae, likely because metamorphosis is stressful itself. Our results highlight that identifying effects of aquatic pollutants and synergisms with natural abiotic stressors, not only in the aquatic larval but also in the terrestrial adult stage, is crucial to fully assess the ecological impact of aquatic pollutants and to reveal the impact on the receiving terrestrial ecosystem through a changed aquatic-terrestrial subsidy.

## 1. Introduction

Our knowledge on the effects of exposure to aquatic pollutants is mainly limited to the aquatic stage of organisms, thereby potentially underestimating their ecological impact (Bundschuh et al., 2022; Henry and Wesner, 2018; Rasmussen et al., 2018). Many organisms have a complex life cycle with aquatic larvae and terrestrial adults where exposure during the aquatic larval stage may cause negative carry-over effects in the unexposed terrestrial adult stage (Stoks and Córdoba-Aguilar., 2012; Stoks et al., 2022). Such delayed effects across metamorphosis are understudied (but see e.g., mayflies: Beketov and Liess, 2005, Wesner et al., 2014, amphibians: Rohr et al., 2017, and damselflies: Bots et al., 2010). This is concerning because some experiments revealed negative effects of exposure to pollutants in the aquatic larval stage to be stronger or even only detectable in the terrestrial adult stage (e.g., Debecker et al., 2017; Henry and Wesner, 2018; Wesner et al., 2014). Especially studies that looked at long-term effects of larval exposure to pollutants on adult fitness components such as adult lifespan in animals with a complex life cycle are rare (but see Debecker et al., 2017).

Another challenge when assessing the impact of pollutants is that natural populations often face co-exposure to pollutants and natural abiotic stressors, which moreover may magnify their toxicity (Liess et al., 2016). While such synergisms are well documented in the aquatic stage, we have limited information on how potential interactive effects with abiotic stressors, such as food limitation, cross metamorphosis and affect long-term fitness components in the adult stage. The few studies on this topic, mainly limited to short-term effects during or directly after metamorphosis, provide mixed results. While there is some evidence that carry-over effects were still interactive in the adult stage (Campero et al., 2008; Janssens and Stoks, 2013) this was not always the case (Fong et al., 2016; Rohr et al., 2004). These studies only considered short-term fitness effects, while long-term effects on adult fitness may give a more complete picture to assess the ecological impact of delayed interactive effects of aquatic pollutants.

One pollutant whose toxicity to aquatic organisms is poorly understood is 2,4-dinitrophenol (DNP) (Kwak et al., 2020). DNP is an industrial chemical used as wood preservative and pesticide (Harris and Corcoran, 1995; Mwesigwa et al., 2000) and has been classified as very toxic for aquatic life by ECHA (2019). Negative effects of DNP exposure on aquatic organisms include increased mortality (Kennedy et al., 2015) and declines in growth and development rates (Salin et al., 2012). DNP acts by uncoupling the mitochondrial oxidative phosphorylation, thereby reducing ATP production (Lou et al., 2007). Therefore, interactive effects between DNP exposure and food limitation are especially relevant to study. Food limitation is common in natural populations (Metcalfe and Monaghan, 2001) and has been shown to magnify the toxic effects of many pollutants (reviewed in Holmstrup et al., 2010; Shahid et al., 2019). Nevertheless, to our knowledge, no other research has examined the combined effect of DNP and chronic food limitation.

The aim of this study was to investigate the impact of aquatic exposure to DNP and how this was modulated by food limitation in the larvae of a semi-aquatic insect, and to test delayed effects during and after metamorphosis on long-term fitness of the terrestrial adults. As study species, we used the damselfly *Lestes viridis*. Damselflies are well-studied semi-aquatic insects that are in their obligate aquatic larval stage vulnerable to pollutants as they cannot escape exposure (Beketov et al., 2013; Stoks et al., 2015). To better understand the single and combined effects of DNP and food limitation on life history traits during the larval (growth rate, survival, development) and adult stage (mass at emergence and adult lifespan), we also measured larval foraging-related behaviours and a set of physiological variables related to the mode-of-action of DNP. Given the expected effects of DNP and food limitation, we quantified the net energy budget by integrating the energy storage and the cellular metabolic rate (De Coen and Janssen, 2003), and the ATP/ADP ratio as an indicator for the metabolic capacity of an organism (Hanski et al., 2004). We also determined the oxidative damage to the lipids by measuring lipid peroxidation (Monaghan et al., 2009). Our main hypotheses were the following: (1) single exposure to DNP and to food limitation will negatively influence the larval life history and physiological variables (Kwak et al., 2020), and (2) cause negative carry-over effects of the larval stressors to the unexposed adult stage (e.g., Beketov and Liess, 2005), (3) the negative effects of exposure to DNP on the larvae will be stronger under food limitation (Holmstrup et al., 2010), and (4) this synergistic effect will bridge metamorphosis and negatively affect adult life history traits.

## 2. Materials and methods

### 2.1. Collection and housing

Twigs containing *L. viridis* eggs were collected in January 2016 at a pond in Edegem (51°09’19.1”N, 4°23’55.6”E) in Belgium. The twigs were placed in trays filled with dechlorinated tap water and were gradually warmed to a final temperature of 20 °C. After hatching, the larvae were placed per ten in plastic cups to enhance survival (De Block & Stoks, 2003). Two weeks later, larvae were transferred to individual cups and kept under standard temperature (20 °C), light (14:10 L:D) and food (ad libitum *Artemia* nauplii) conditions for one more week until the experiment started.

### 2.2. Experimental setup

To test for the single and combined impact of DNP exposure and food stress, we set up a full factorial experiment where larvae were exposed to one of the four combinations of two DNP treatments (absent versus present) and two food levels (high versus low). The treatments started three weeks after egg hatching and were continued during the entire larval stage. Response variables during the larval stage were mainly quantified in the final instar. This is motivated by the following reasons: (i) larvae in this instar have been exposed for the longest time to the stressors, hence any effects would be more obvious, (ii) this instar has the longest duration and here the biggest increase in larval mass occurs, and (iii) this instar is closest to the adult stage, hence most relevant when studying carry-over effects into the adult stage. We started 400 larvae per DNP treatment at high food and 500 larvae per DNP treatment at low food, resulting in 1800 larvae at the start of the experiment.

Larvae from the high food level received ad libitum *Artemia* nauplii six days a week. After moulting into the final instar, larvae additionally were fed three chironomids three times a week (based on De Block and Stoks, 2005). Larvae from the low food level received the same amount of *Artemia* three times a week, and in the final instar additionally were fed one chironomid three times a week.

For larvae exposed to the pesticide we used a concentration of 2 mg/L DNP. This concentration of DNP is within the range found in natural water bodies where concentrations up to 12 mg/L have been reported (Nabais et al., 2007). This concentration is expected to be harmful given it is near the chronic HC50 concentration of 2.07 mg/L DNP based on six freshwater species (Kwak et al., 2020). We prepared a stock solution of 20 mg/mL DNP in dimethyl sulfoxide (DMSO) which was stored in the dark at room temperature. This solution was diluted ten times with Milli-Q water resulting in a concentration of 2 mg/mL; from this mixture 100 μL was added to the cups (which were filled with 100 mL of dechlorinated tap water), resulting in a concentration of 2 mg/L. We added the same amount of DMSO to the control conditions. This concentration of DMSO was not harmful for the animals. We placed the larvae in a new cup with DNP (or DMSO for control larvae) when larvae reached their final instar, this was ∼ 52 days after the DNP exposure started. Given the half-life of DNP is ∼ 68 days in aerobic water (Capel and Larson, 1995), this ensured continuous exposure to DNP throughout the larval stage as is realistic in edge-to-field water bodies.

### 2.3. Life history and behavioural response variables

Larval mortality and moults into the final instar were checked three times a week. Based on the latter we calculated the time needed to reach the final instar (development time until F0). We quantified growth rate during the first seven days of the final instar by weighing the animals one day after moulting and seven days later using an electronic balance (Mettler Toledo® AB135-S) to the nearest 0.01 mg. Growth rate was calculated as (ln_final mass_ – ln_initial mass_) / 7 days (McPeek et al., 2001) on 304 – 400 animals per treatment combination. At day 7 in the final larval instar a subset of the larvae (25 – 26 per treatment combination) was used to score foraging-related behaviours (based on Janssens and Stoks, 2012). We placed an individual larva in a plastic container (11 × 11 × 6 cm), which was filled with 100 mL dechlorinated tap water. The larvae were given 10 minutes to acclimate in the container before the observations started. Then, we added 1 mL of ad libitum *Artemia* nauplii to the container, where after we recorded the number of feeding bites, head orientations toward the prey and walks during seven minutes. Another subset of larvae (9 – 13 per treatment combination for all physiological analyses, except for the ATP/ADP ratio that had 13 – 28 larvae per treatment combination) was stored at -80 °C for physiological analyses (see below).

We daily checked for emergence of new adults. The development time was calculated as the total duration of the larval stage. We estimated mortality during metamorphosis as the percentage of animals that died during metamorphosis relative to the numbers that survived up to metamorphosis. Metamorphosis success was scored using three categories (Dinh et al., 2016): 0 = animal that did not fully cast its larval skin and died during metamorphosis, 1 = animal that fully casted its skin, but showed malformations of abdomen and/or wings, and 2 = animal that fully casted its skin and had no malformations. One day after emergence, the successfully emerged adults (category 2) were weighed until the nearest 0.01 mg and the sex was determined. A subset of adults (4 – 10 per treatment combination for all physiological analyses, except for the ATP/ADP ratio that had 5 – 26 adults per treatment combination) was directly frozen (−80 °C) for physiological analyses. To determine adult lifespan, the other adults (59 – 142 per treatment combination) were individually marked on their wing with a permanent marker and placed in insectaries (30 × 30 × 30 cm) covered with netting that contained *ad libitum* fruit flies as food source. Males and females were housed in separate insectaries with a maximum of ten individuals per insectary. We daily monitored survival of the adults.

### 2.4. Physiological response variables

A set of physiological variables was quantified on the supernatants of both larval and adult bodies using spectrophotometry. For all the physiological variables except for the ATP/ADP ratio analyses, we homogenized the bodies and diluted them seven times using phosphate buffer saline (PBS, 50mM, pH 7.4), centrifuged the mixture for 5 minutes (16,100 g, 4 °C) and used the supernatant for the physiological analyses.

To determine the cellular energy allocation (CEA) or total net energy budget of each larva we divided the energy available (Ea) by the energy consumed (Ec) (see e.g., Pestana et al., 2009; van Dievel et al., 2019; Verslycke et al., 2004). To calculate the Ea, we summed the amount of energy stored in the three main energy reserve biomolecules (fat, sugars and proteins expressed in mg per mg body mass). These bio-energetic variables, are widely used sensitive biomarkers to stressors such as toxicants, advocated, amongst others by De Coen & Janssen (2003). With the here used assay we indeed have shown before in damselfly larvae, that stressors such as pollutants and warming may negatively affect these bio-energetic variables, and moreover, we could demonstrate strong links among stressor treatment groups between the levels of CEA and life history traits (survival and growth rate), indicating that stressors (partly) affect life history through their effects on these bio-energetic variables (see e.g. Verheyen and Stoks 2021). After spectrophotometric quantification of these biomolecules (see Appendix S1 and S2 for details), their concentrations were converted into their energetic equivalents using the following energy of combustion values: 39,500 mJ/mg for lipids, 17,500 mJ/mg for sugars, and 24,000 mJ/mg for proteins. The Ec was determined by measuring the activity of the electron transport system (ETS) which is a proxy for the cellular metabolic rate (De Coen and Janssen, 2003). The multi-enzyme ETS complex is situated in the mitochondria’s inner membrane where it serves as a link between oxygen and the organic matter that is being oxidized (G.-Toth et al., 1995). Specifically, for an average mixture of lipids, sugars, and proteins we used an oxyenthalpic equivalent of 484 kJ/mol O_2_ to convert the total oxygen consumed to energetic equivalents (based on De Coen and Janssen, 2003).

As a proxy for oxidative damage to lipids we quantified the concentration of malondialdehyde (MDA) using the optimized assay of the thiobarbituric acid assay (TBA assay) following the protocol described in Janssens and Stoks (2017) which was based on (Miyamoto et al., 2011). Using this assay, we have shown before single and interactive effects of pollutants and natural stressors in damselfly larvae (e.g. Janssens & Stoks 2017). We filled a microtube with 50 μL TBA solution (0.4 %, in 0.1 M HCl) and 50 μL of the supernatant and incubated the mixture at 90 °C for 60 minutes. Afterwards, we added 165 μL butanol, and mixed and centrifuged the tubes for 4 minutes at 3000 rpm. From the upper layer we used 30 μL to fill the wells of a 384 well microtiter plate and measured fluorescence at an excitation/emission wavelength of 535/550 nm. We calculated the concentration of MDA based on a standard curve of 1,1,3,3-tetramethoxypropan 99 % malonaldehyde bis (dimethyl acetol) 99 %. Measurements were done in duplicate, and the means were used for the statistical analyses. MDA concentrations were expressed as nmol per mg fat.

We quantified the ATP/ADP ratio based on a modified version of the protocol from Liu et al. (2006). Using this assay, effects of stressors such as exposure to oil on this ATP/ADP ratio were shown (Li et al. 2017). First, we homogenized the bodies of the animals in perchloric acid (PCA 7 %) (body mass x 15 μL). After putting the mixture on ice for 1 hour, we added 93 μL neutralization solution (3M KOH + 3M KCl + 0.4 M Tris) to 250 μL of the mixture. After thorough mixing, we centrifuged the tubes at 10,000 rpm for 5 minutes at 4 °C. From the supernatant, 250 μL was brought into an Eppendorf tube with filter and centrifuged at 13,000 g for 2 minutes at room temperature. We used 20 μL of the filtered solution for high-performance liquid chromatography (HPLC) (DGU-20A3, Shimadzu, Japan). For HPLC, we used a C18 column (250 × 4.6 mm, 5 μm), a flow rate of 1 mL/minute, a 20 μL loop and a gradient of two solutions: solution A (60 mM K_2_HPO_4_ + 40 mM KHPO_4_, pH 6.65) and solution B (acetonitrile + 0.1 % trifluoroacetic acid). The amounts of ATP, ADP and AMP were calculated based on standard curves of these molecules. Following, for example, Yang et al. (2014), we report data on the ATP/ADP ratio; these patterns were similar as for the ATP/(ADP+AMP) ratio (results not shown).

### 2.5. Statistical analyses

All statistical analyses were done in Rstudio v1.4.1103 (Rstudio Team, 2021). All the plots were made using the ggplot package for R (version: 3.3.3; Wickham, 2016). Survival (binary) was analyzed with a generalized linear model with a quasi-binomial error distribution and logit link. The metamorphosis success (0, 1 or 2) was analyzed with an ordered multinomial logistic regression model. Development time, the behavioural variables (feeding strikes, head orientations and walks) and adult lifespan were analyzed with Poisson regression models with a log-link. The behavioural variables were corrected for larval mass, by adding larval final mass as a covariate in these models. All other life history (growth rate and mass at emergence) and physiological variables were analyzed using linear effect models. We analyzed the physiological variables separately per life stage, given the strong differences in the measured trait values between life stages. Food level, DNP exposure and their interaction were added as independent variables in all the models. In both the larval and adult data sets of the variable ATP/ADP, one outlier was removed. For the larvae, ATP/ADP was boxcox-transformed to meet the assumptions of normality of the residuals and homogeneity of variances. In all the models, the ‘car package’ was used to calculate the F-statistics as well as the p-values. Pairwise contrasts, utilizing estimated marginal means with FDR correction, were used to further analyze the significant interactions (emmeans package: version 19 1.5.4; Lenth, 2021). The full statistical tables and figures of the physiological variables can be found in the Appendix S5.

The type of interaction (additive, antagonistic or synergistic) between the stressors DNP and food limitation (low food) was formally determined for the variables larval survival, metamorphosis success and adult lifespan. This was done because for survival-related variables general linear models the test for interaction patterns can be misleading as an animal killed by one stressor cannot be killed by a second stressor (Schäfer and Piggott, 2018). Therefore, for these variables we used the independent action (IA) model, where a synergistic interaction was concluded if the predicted combined effect was larger than the upper limit of the confidence interval 95 % confidence interval of the observed combined effect (Coors & de Meester, 2008). As a measure of the strength of a synergism we quantified the model deviation ratio (following Shadid et al., 2019). The higher the model deviation ration above 1 (with 1 indicating an additive effect), the stronger the synergism. In the appendix S6, more details can be found about the application of the IA model as well as the results.

## 3. Results

### 3.1. Life history and behavioural response variables

Both exposure to DNP and to the low food level reduced survival until adult emergence (Table 1, Fig. 1A). While the GLM detected no significant interaction between both stressors, DNP decreased the survival ∼ 61 % at low food and ∼ 39 % at high food. The stronger DNP-imposed mortality at low food was confirmed by the independent action model identifying the interaction between DNP and low food as synergistic with a model deviation ratio of 1.86 (see Appendix S6). Exposure to DNP and the low food level each reduced the larval growth rate and increased the larval development time (Table 1, Fig. 1B-C). In addition, the DNP-imposed growth reduction was significantly larger at high food (∼ -16 %) than at low food (∼ -14 %) (DNP × food level interaction, Table 1), yet this subtle 2 % difference is unlikely to be biologically relevant.

**Table 1.** Results of (G)LMs testing for the effects of DNP exposure and food level on the life history variables of the damselfly *Lestes viridis*.

| Effect | Larval survival |  |  | Larval growth rate |  |  | Larval development time |  |  |
| --- | --- | --- | --- | --- | --- | --- | --- | --- | --- |
| | df | $\chi^2$ | p | df<sub>1</sub>,df<sub>2</sub> | F | P | df | $\chi^2$ | p |
| DNP | 1 | 476.06 | <0.0001 | 1,1369 | 48.91 | <0.0001 | 1 | 7.30 | 0.0069 |
| Food level | 1 | 51.72 | <0.0001 | 1,1369 | 464.35 | <0.0001 | 1 | 962.01 | <0.0001 |
| DNP x Food level | 1 | 1.77 | 0.18 | 1,1369 | 5.97 | 0.015 | 1 | 0.12 | 0.73 |
|  | Metamorphosis success |  |  | Mass at emergence |  |  | Adult lifespan |  |  |
| | df | $\chi^2$ | p | df<sub>1</sub>,df<sub>2</sub> | F | P | df | $\chi^2$ | p |
| DNP | 2 | 71.31 | <0.0001 | 1,540 | 2.03 | 0.16 | 1 | 1787.81 | <0.0001 |
| Food level | 2 | 222.75 | <0.0001 | 1,540 | 195.87 | <0.0001 | 1 | 1154.64 | <0.0001 |
| DNP x Food level | 2 | 3.61 | 0.16 | 1,540 | 3.36 | 0.067 | 1 | 893.96 | <0.0001 |

**Figure 1.**
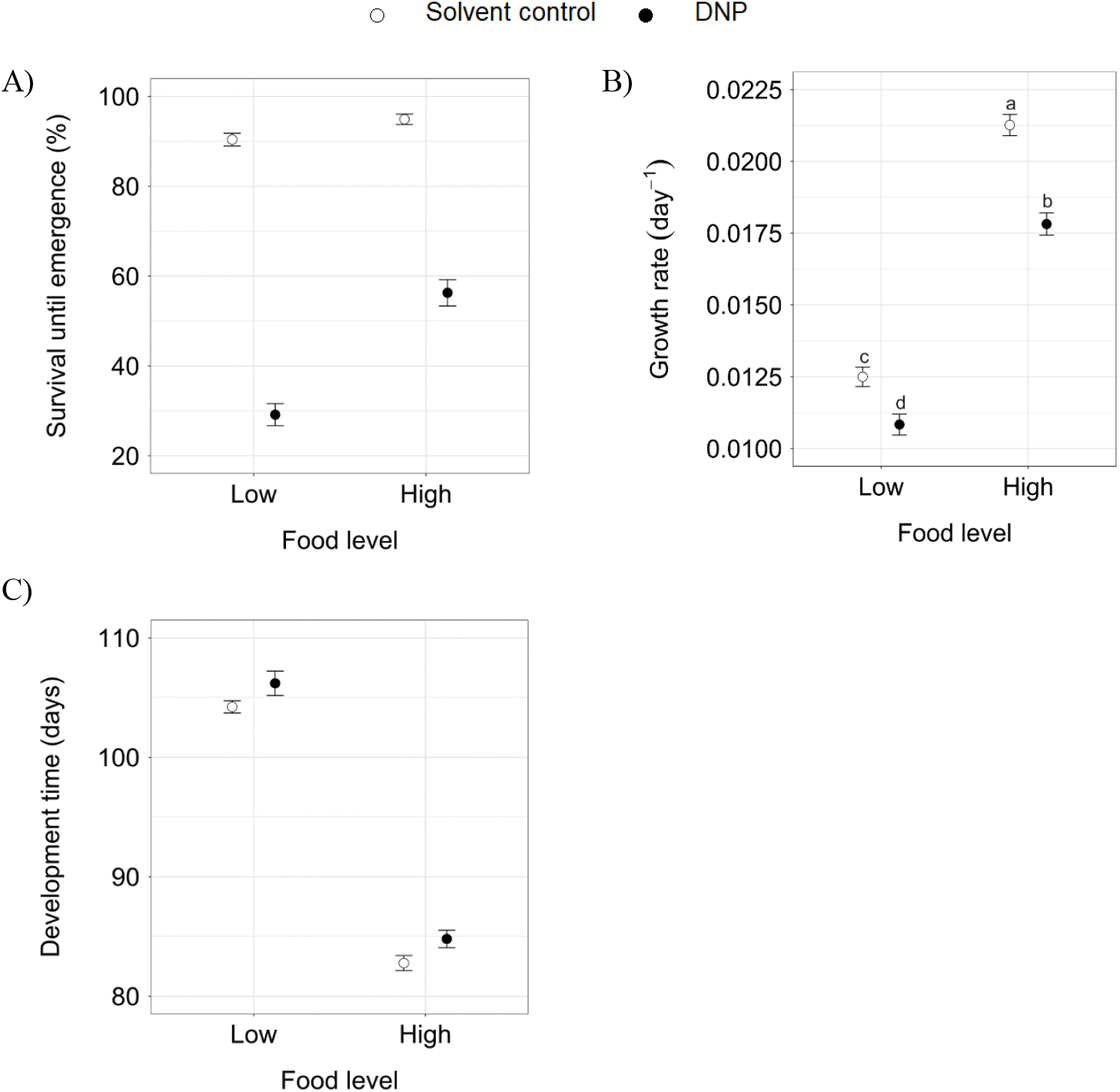
The effect of DNP exposure and food level on the larval life history variables of the damselfly *Lestes viridis*: (A) survival until emergence, (B) growth rate and (C) development time. The means ± 1 SE are given. Different letters denote significant differences based on pairwise contrasts from the significant DNP × food level interaction.

Both exposure to DNP and to low food increased mortality during metamorphosis and reduced successful emergence, i.e., emergence with normal wings (Table 1, Fig. 2). Although, the interaction between DNP and food level was not significant, the independent action model identified a synergistic interaction between the stressors with a model deviation ratio of 2.27 (see Appendix S6). Exposure to DNP increased mortality during metamorphosis more at low food (∼ 38 %) than at high food (∼ 13 %). Furthermore, DNP reduced successful emergence more at low food (∼ 35 %) than at high food (∼ 16 %). From those adults that emerged successfully, low food during the larval stage decreased the mass at emergence, while DNP had no main effect (Table 1, Fig. 3A). There was a trend (p = 0.067) for a DNP × food level interaction indicating that DNP had no effect at low food whereas DNP caused a mass reduction of ∼3 % at high food.

**Figure 2.**
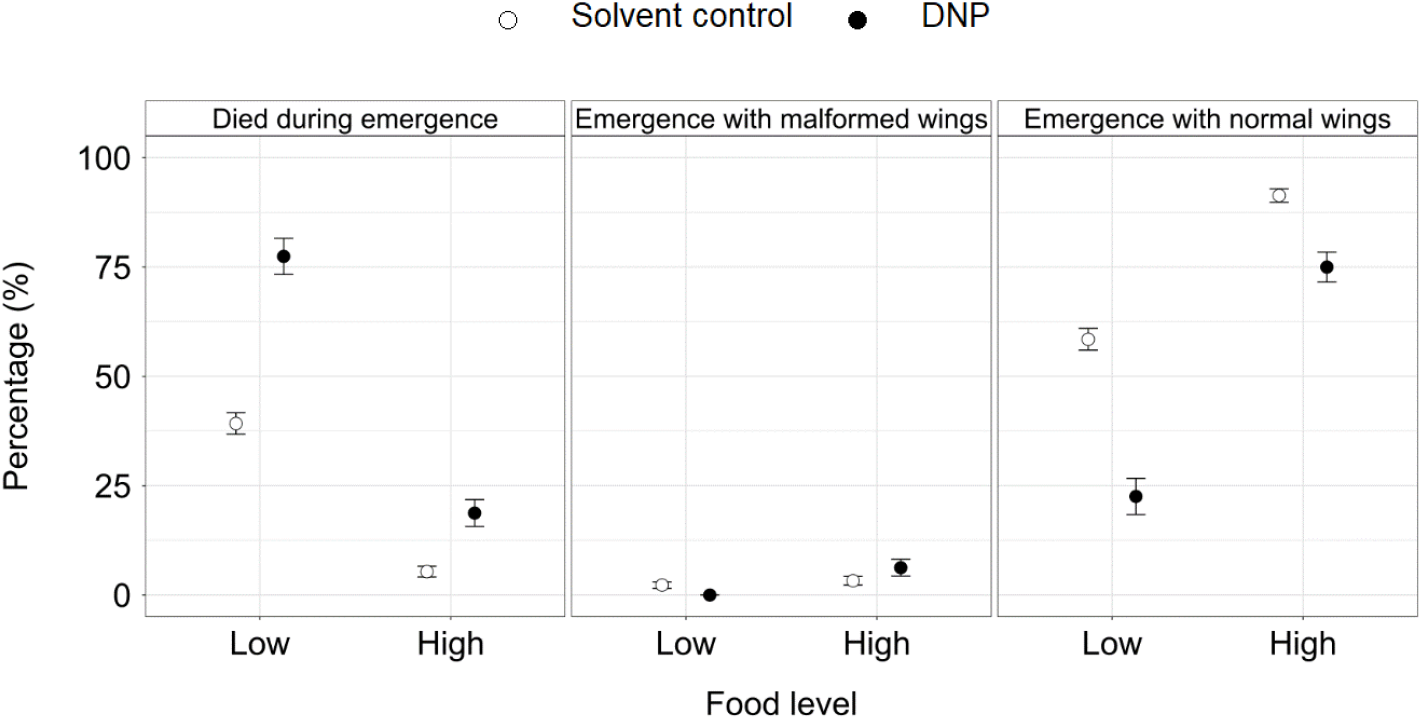
The effect of DNP exposure and food level on the metamorphosis success of the damselfly *Lestes viridis*. The means ± 1 SE are given.

**Figure 3.**
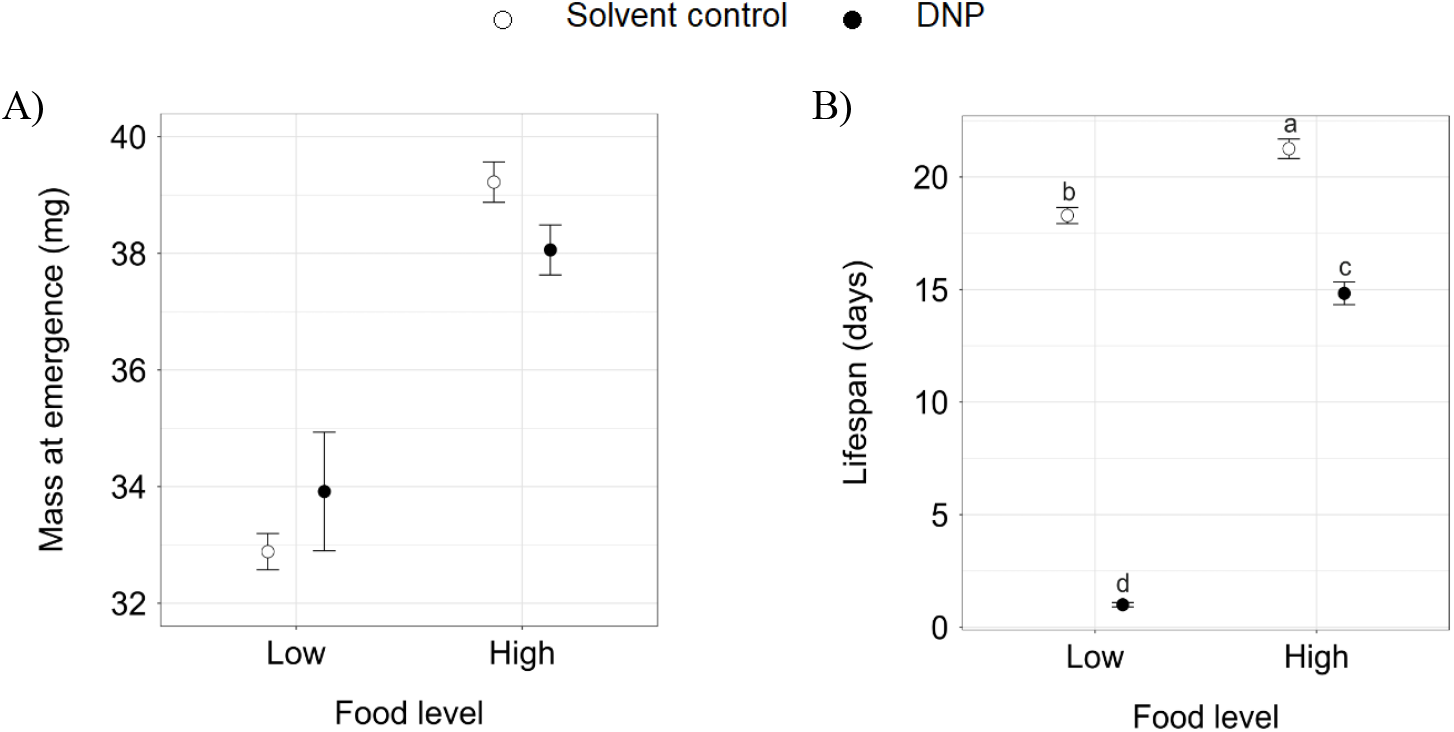
The effect of DNP exposure and food level on the adult life history variables of the damselfly *Lestes viridis*: (A) mass at emergence and (B) adult lifespan. The means ± 1 SE are given. Different letters denote significant differences based on pairwise contrasts from the significant DNP × food level interaction.

DNP exposure as well as low food decreased the adult lifespan (Table 1, Fig 3B). The DNP-imposed lifespan reduction was larger at low food (∼ -95 %), where all the adults died within 1 day after emergence, than at high food (∼ -31 %) (DNP × food level, Table 1). Accordingly, the interaction between DNP and low food was identified as highly synergistic based on the independent action model with a model deviation ratio of 12.80 (see Appendix S6).

DNP exposure caused a ∼30 % reduction in all three mass-corrected behavioural activities (the number of feeding strikes, head orientations and walks), but only at high food and not at low food (DNP × Food level, Table S4.1, Fig. S4.1). The food level itself had no effect in the absence of DNP.

### 3.2. Physiological response variables

In the larvae, DNP exposure did not affect the available energy (Ea) and the consumed energy (Ec), whereas the low food level decreased both Ea (∼ -20 %) (main effect Food level: F_1,41_ = 18.02, p = 0.0001) and Ec (∼ -17 %) (main effect Food level: F_1,41_ = 5.13, p = 0.03) (Table S5.1, Fig. S5.1A-B, see details on components of Ea in Appendix S3). Neither DNP nor the food level influenced the cellular energy allocation (CEA) (Table S5.1, Fig. S5.1C). DNP exposure had no effect on the MDA levels, but lower food resulted in decreased MDA levels (∼ -47 %) (main effect Food level: F_1,41_ = 9.20, p = 0.0042) (Table S5.1, Fig. S5.1D). Neither DNP nor low food affected the ATP/ADP ratio (Table S5.1, Fig. S5.1E).

In the adults, Ea was not affected by either the DNP or food treatments experienced as larva (for the components of Ea, see Appendix S3). Ec was only reduced by DNP at high food (contrast: p = 0.025) but not at low food (contrast: p = 0.90), despite the non-significant DNP × food level interaction (Table S5.2, Fig. S5.2A-B). DNP had no effect on the CEA, but low food increased the CEA (∼ 15 %) (main effect Food level: F_1,25_ = 4.58, p = 0.042) (Table S5.2, Fig. S5.2C). Neither DNP nor food level influenced the MDA levels or the ATP/ADP ratio (Table S5.2, Fig S5.2D-E).

## 4. Discussion

We investigated the impact of chronic exposure to DNP and how this was modulated by limited food availability in the larval stage, and the possible delayed effects during and after metamorphosis in the adult stage of the damselfly *L. viridis*. The applied ecologically relevant DNP concentration and the low food level imposed in the larval stage, each had large negative effects on the life history, behaviour and to a lesser extent on the physiology of not only the larvae but also the adults. As expected, the negative impact of DNP was more pronounced at low food as illustrated by stronger negative effects on larval survival, metamorphosis success and especially on adult lifespan. These stronger negative effects reflect a synergism for these survival-related variables between both stressors.

### 4.1. The effects of DNP at high food

All studied larval and adult life history traits were negatively affected by DNP exposure at high food indicating the here tested DNP concentration to be stressful: DNP decreased larval survival and growth rate, increased development time and decreased mass at emergence and adult lifespan. These effects on life history were associated with an overall decreased larval activity. In sharp contrast, the physiological variables were unaffected by DNP exposure, except for an effect on energy consumption in the adult stage. The assays of the bio-energetic variables (Ea, Ec and CEA), and the proxy for lipid peroxidation (MDA) have proven to possess enough resolving power to detect impacts of pollutants and natural stressors in damselfly larvae (e.g. Janssens and Stoks 2017, Verheyen and Stoks 2021). Nevertheless, we note that the observed ranges of values of several variables in our study were rather narrow. Therefore, we cannot fully exclude that some smaller effects went undetected.

Negative effects of the applied DNP concentration (2 mg/L) on life history were expected given it is near the chronic HC_50_ concentration of 2.07 mg/L DNP based on six freshwater species (Kwak et al., 2020). In line with our results, the relative few studies on freshwater animals showed DNP, for example, to decrease embryo survival in the fish *Oryzias latipes* (Kwak et al., 2020) and juvenile survival in the water flea *Ceriodaphnia dubia* (Kennedy et al., 2015). Several non-exclusive and potentially interacting mechanisms may explain these negative effects on life history. We list them here, and make clear for which mechanism there is explicit support for the link with life history, but also note that these studies used other assays to measure the physiological/biomolecular variables. First, DNP can reduce the lysosomal stability (Fent and Hunn, 1996), which causes the release of lysosomal hydrolases in the cytoplasm, leading to cell death (Boya, 2012). Lysosomal destabilization can therefore be a potential toxicity mechanism (Zhao et al., 2021). Second, DNP can inhibit the activity of the enzyme succinate dehydrogenase in the mitochondria (Fent and Hunn, 1996), and thereby disrupt the electron flow to the respiratory chain ubiquinone pool, resulting in oxidative stress (Rustin et al., 2002). Third, DNP may cause protons to flow over the plasma membrane, thereby depolarizing it, which can lead to negative effects on physiological functions (Jastroch et al., 2014). Fourth, DNP exposure is expected to reduce the ATP production (Lou et al., 2007), which may result in less energy available to invest in growth and development. This mechanism was demonstrated to underlie negative effects of DNP on growth and development in *Rana temporaria* tadpoles (Salin et al., 2012). Yet, our results do not seem to support this mechanism as we did not observe a reduced ATP/ADP ratio nor a reduced net energy budget (CEA). We do note, however, that none of our treatments significantly affected the ATP/ADP ratio, and only the food level had a significant effect on CEA, hence we cannot exclude our assays were not able to detect small effects. Fifth, the observed negative effects may also be the result of an increased investment in mechanisms to compensate for the expected reduced ATP production per mitochondrion under DNP exposure. Three such compensatory mechanisms have been described: increasing the capacity of the oxidative phosphorylation in each of the mitochondria, increasing the biogenesis of the mitochondria and/or activating the glycolysis (Salin et al., 2012; Schlagowski et al., 2014). These compensatory mechanisms may be one reason why the net energy budget and the ATP/ADP ratio were not reduced as could have been expected under a lower mitochondrial ATP production (Geelen et al., 2008; Sibille et al., 1995). Finally, the observed reduction in foraging activity may also have negatively affected life history as foraging activity is tightly linked to growth rate in damselfly larvae (Brodin and Johansson, 2004; Stoks et al., 2012). A similar reduction in larval activity under DNP exposure was also documented in, for example, *Danio rerio* zebra fish, where it was explained by the lower ATP levels resulting in insufficient energy for muscle movement (Marit and Weber, 2011). Our results do not seem to support this explanation as we could not detect an effect on ATP levels, but again we cannot exclude the assay was not able to detect small effects. In any case, several of the other discussed mechanisms could explain the reduced foraging activity in our study.

Notably, chronic DNP exposure during the larval stage had delayed costs during metamorphosis (increased mortality) and after metamorphosis (lowered mass at emergence and shortened adult lifespan). Chronic DNP exposure from the juvenile stage onwards has been shown to shorten the adult lifespan in zebra finch (Stier et al., 2021), yet, and in contrast with our study, not only the juvenile but also the adult zebra finches were exposed to DNP. Similar to current findings, in a previous study on *L. viridis* where all larvae received high food, chronic exposure to DNP also seemed to reduce the adult lifespan, but this effect did not reach statistical significance, likely because of low sample sizes (Janssens and Stoks, 2018). Our current results add to the accumulating evidence that larval exposure to pollutants may negatively affect adult fitness (e.g., Beketov and Liess, 2005; Bots et al., 2010; Van Dinh et al., 2016; Rohr et al., 2017). Such carry-over effects of larval stressors across metamorphosis are still poorly understood (Stoks et al., 2022), but crucial to document when assessing the total impact of pollutants on organisms.

The energy consumption (measured as the cellular metabolic rate) was not affected by DNP in the larvae, yet was decreased in the adults. This is in contrast with other studies demonstrating that DNP increased the metabolic rate (Caldeira Da Silva et al., 2008; Salin et al., 2012; Stier et al., 2021, 2014). The latter pattern was explained by a reduced efficiency of the oxidative phosphorylation in the mitochondria, whereby more oxygen needs to be consumed for a given ATP amount, thereby increasing the metabolic rate (Salin et al., 2012). We hypothesize that larvae did not detectably increase their metabolic rate to save energy to invest in defense mechanisms. Intriguingly, while no effect was observed in DNP-exposed individuals during the larval stage, they showed a decreased metabolic rate in the adult stage. Effects of a stressor on a trait may only become visible in the adult stage as metamorphosis itself is a stressful event (Campero et al., 2008) that may cause, for example, oxidative stress (Gaupale et al., 2012). Together with the DNP exposure in the larval stage this may have caused the adults to reduce their metabolic rate (cfr metabolic depression, Storey, 2015; for damselfly larvae: Van Dinh et al., 2016).

### 4.2. The effects of low food in the absence of DNP

As expected, low food in the larval stage negatively affected all studied larval and adult life history traits: it decreased larval survival and growth rate, increased the development time, and decreased the adult mass and lifespan. These are general life history costs that typically are attributed to a lower energy content (e.g., Beketov and Liess, 2005; De Block and Stoks, 2008; Janssens et al., 2014) as indeed was observed in the larvae. While larvae reared at the low food level likely had a lower food intake during development, this was not reflected in the behavioural trials. Note, however, all larvae were given the same amount of food during the behavioural trials.

Despite the likely lower accumulated food intake throughout the larval stage, the net energy budget (CEA) was not reduced in the larvae, because they also lowered the energy consumed (metabolic rate) under food stress (as previously documented in the study species: Stoks et al., 2006). On its turn, the latter may explain the decrease in oxidative damage (MDA). In the adult stage, the energy consumed was also lower when reared as larvae at low food, yet the available energy was not affected, resulting in an increased net energy budget. Given that the low food level reduced the energy storage in the larvae and as the adult physiology was measured 24h after emergence, when the adults had not eaten yet, the absence of an effect of low food on the adults’ energy storage is surprising. While we cannot exclude that our assays may not have detected small effects of low food on Ea, Probably, the absence of a clear effect is due to strong survival selection. Indeed, a considerably percentage (∼ 41 %) of the larvae reared at low food did not metamorphose successfully. Likely these larvae that died, hence could not be measured as adults, were those with the lowest energy reserves.

### 4.3. The effects of DNP in the presence of low food

DNP had stronger negative effects on the larval survival, survival during metamorphosis and on the adult lifespan when combined with low food; this reflects synergistic interactions between both stressors. In contrast, we could not detect any negative effects of DNP on the overall behaviour of the larvae at low food while such effect was detected at high food. Possibly, these energy-limited larvae faced a stronger pressure to obtain energy for defense and repair when exposed to DNP compared to larvae reared at high food. Alternatively, the higher DNP-caused larval mortality at low food than at high food, may have removed the larvae with the lowest food intake from the dataset.

The synergisms between DNP and low food that were detected for the three survival-related variables that capture mortality during the larval stage, during metamorphosis and during the adult stage, had a strong total impact on survival. Our study thereby provides a rare example of interactive carry-over effects between larval exposure to a pollutant and a natural stressor on long-term adult fitness (see also Debecker et al., 2017). These synergisms can be due to the observed lower energy content at low food, hence a lower ability to compensate and defend against the negative influence of DNP and are expected to occur in a wide range of pollutants (Sibly and Calow, 1989). In line with this, many studies have shown pollutants to be more toxic at lower food levels (e.g., pesticides: Shahid et al., 2019).

A striking finding was that the observed synergism between DNP exposure and food stress in the larval stage became stronger when larvae metamorphosed and was especially strong in the adults. Indeed, all adults that emerged after being exposed as larvae to DNP at low food died within 24h after emergence. This illustrates the importance of quantifying synergisms with aquatic pollutants not only during the larval stage, but also during and especially after metamorphosis. Given that the metamorphosis process itself is stressful and metabolically costly (Gaupale et al., 2012; Lowe et al., 2021), this likely left the adults that were food-limited as larvae with not enough energy to protect themselves against the negative influence of DNP. An alternative mechanism through DNP bio-accumulation causing stronger DNP effects later in life seems less likely given the low bio-accumulation potential of DNP (ECHA, 2019; USEPA, 2015).

## 5. Conclusions

Our results provide important novel information for the much needed improvement of risk assessment of aquatic pollutants in the many species with aquatic life stages and terrestrial adults (Henry and Wesner, 2018; Rasmussen et al., 2018). Our study adds to the few others demonstrating that exposure to pollutants in the larval stage can on itself and through interactive effects with natural stressors affect the long-term fitness of the terrestrial adults. Notably, and at first sight counterintuitive, our results for survival-related traits illustrate that such synergistic effects between stressors can be stronger during the unexposed adult stage than during the exposed larval stage. Given this may be explained by the stressful metamorphosis process (Lowe et al., 2021), we hypothesize that such pattern of stronger synergisms after metamorphosis to be widespread. Identifying effects not only in the aquatic life stages but also in the terrestrial adult stages is important to fully assess the impact of aquatic pollutants on the receiving terrestrial ecosystem (Henry and Wesner, 2018; Bundschuh et al., 2022). Semi-aquatic insects such as damselflies provide an important aquatic-terrestrial subsidy both in terms of food quantity and food quality (Popova et al., 2017). Our results indicate that both aquatic pollutants (e.g., DNP) and natural stressors (e.g., food limitation), can strongly reduce this subsidy, and especially so when both stressor types are combined.

## Acknowledgments

We thank Rony Van Aerschot and Geert Neyens for technical support during the experiment and Ria Van Houdt for performing the physiological analyses. LJ and JV received a postdoctoral fellowship from the Fund for Scientific Research-Flanders (FWO). We benefited from a research grant from the KU Leuven (C16/17/002).

## Abbreviations

ADP: adenosine diphosphate
AMP: adenosine monophosphate
ATP: adenosine triphosphate
CEA: cellular energy allocation
DMSO: dimethyl sulfoxide
DNP: 2,4-dinitrophenol
Ea: energy availability
Ec: energy consumption
ECHA: European Chemicals Agency
ETS: electron transport system
FDR: false discovery rate
(G)LM: (generalized) linear model
HC: hazard concentration
HPLC: high-performance liquid chromatography
IA: independent action model
MDA: malondialdehyde
PCA: perchloric acid
ROS: reactive oxygen species
TBA: thiobarbituric acid
USEPA: United States Environmental Protection Agency

---

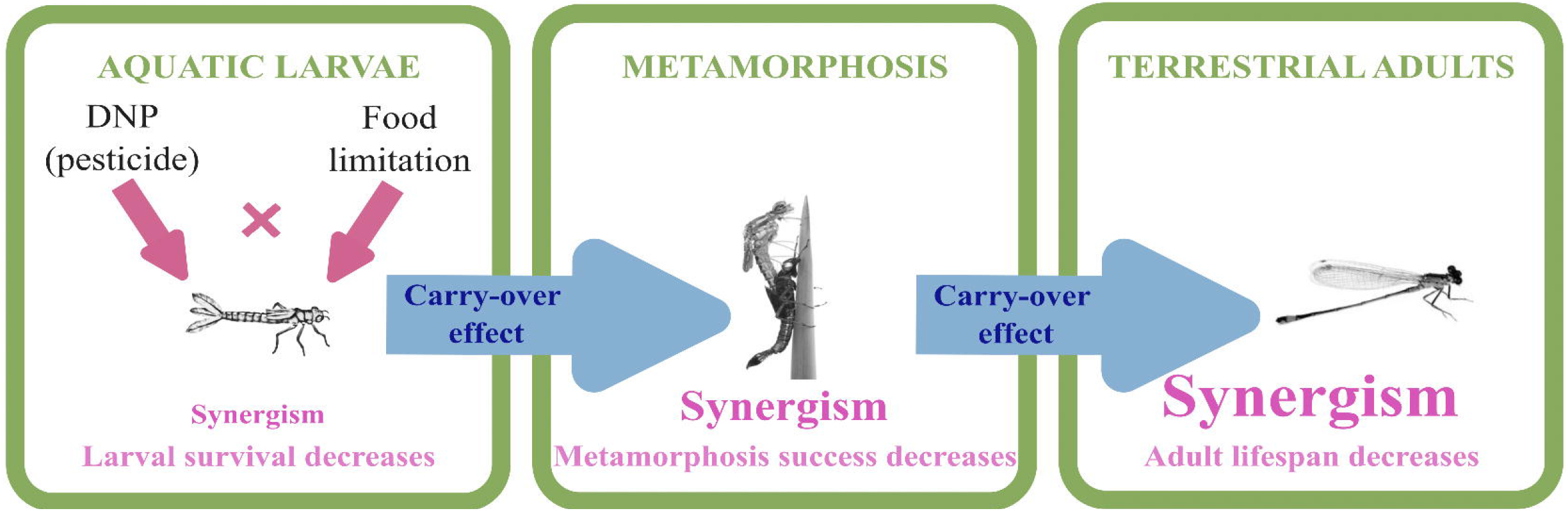

