## Appendix for "Synergistic survival-related effects of larval exposure to an aquatic pollutant and food stress get stronger during and especially after metamorphosis and shape fitness of terrestrial adults"

S1 contains detailed information about the protocol for determining the bioenergetic variables.

S2 contains detailed information about the statistical analyses of the bioenergetic variables.

S3 contains the results of the bioenergetic variables.

S4 contains the full statistical tables and figures of the physiological variables.

S5 contains information about the application of the independent action model for the survival-related variables

The supporting information contains five figures (S3, S4 and S5), six tables (S3, S4, S5 and S6) and is in total 20 pages.

**Appendix S1: Protocol for determining the bioenergetic response variables**

To determine the bioenergetic response variables, we first homogenized the bodies of larvae or adults using a pestle. Second, it was diluted seven times using phosphate buffer saline (PBS, 50mM, pH 7.4) and centrifuged for 5 minutes (16,100 g, 4 °C) whereafter the supernatant was used.

To quantify fat content, we followed the protocol described in Janssens & Stoks, 2014. We filled a glass tube with 8 µL supernatant, and 56 µL sulfuric acid (100 %). After incubating the tubes for 20 minutes at 150 °C, we added 64 µL of Milli-Q water. From this mixture we transferred 30 µL to a 384 well microtiter plate. Absorbance was measured at 340 nm and the fat concentrations were calculated using a standard curve of glyceryl tripalmitate. Measurements were done in triplicate and the means per larva were used for statistical analyses.

We quantified the sugar content (glucose and glycogen: Hahn & Denlinger, 2007), following the protocol described in (Stoks et al., 2006) and worked with the glucose kit of Sigma Aldrich USA. We filled a 96 well microtiter plate with 37.5 µL Milli-Q water, 12.5 µL supernatant and 100 µL glucose assay reagent (Sigma G3293). The samples were incubated for 20 minutes at 30 °C, after which we measured absorbance at 340 nm. Glucose concentrations were calculated based on a standard curve of glucose. Measurements were done in duplicate and the means per larva were used for statistical analyses.

Protein content in the body homogenates was measured using the Bradford method (Bradford, 1976). We mixed 160 µL Milli-Q water with 1 µL supernatant whereafter 40 µL BioRad reagent was added. After further mixing, the absorbance was measured at 595 nm. By using a standard curve of Bovine Serume Albumine, we calculated the protein concentrations. Measurements were done in quadruplicate and the means per larva were used for statistical analyses. Fat, total sugar and protein contents (in mg) were expressed per mg larva.

The quantification of the ETS activity was based on the protocol described in Janssens et al., 2015. We filled the wells of a 384 well microtiter plate with 15 µL buffered substrate solution (0.13 M Tris HCl, 0.3 % Triton X-100, 1.7 mM NADH, 250 µM NADPH, pH 8.5) and 5 µL of the supernatant. Afterwards, we added 10 µL INT (8 mM p-iodonitrotetrazolium) to replace the electron acceptor $O_2$ and furthermore to receive electrons, via NADH-cytochrome oxidoreductase, from NADPH which will lead to the formation of formazan. We immediately followed the increase in formazan absorbance at 490 nm (Infinite M2000, TECAN) and 20 °C during 5 minutes with readings every 20 seconds. The Lambert-Beer law as well as the molecular extinction coefficient (15.9 $mM^{-1}cm^{-1}$) were used to calculate the formazan concentrations. Thereafter, the cellular oxygen consumption rates were determined taken into account that in order to form 2 µmol formazon in the ETS complex, 1 µmol $O_2$ is used. The activity of ETS was indicated as nmol $O_2$ per minute per mg larva.

**Appendix S2: Statistical analysis for determining the bioenergetic response variables**

The statistical analyses were done in Rstudio version 1.4.1103 (Rstudio Team, 2021). All the plots were made using the ggplot package for R (version: 3.3.3; Wickham, 2016). We analyzed the bioenergetic response variables separately per life stage, given the often strong differences in the measured trait values between life stages. In all the models food level, DNP exposure and their interaction were added as independent variables. We used generalized linear models (LMs) for all the variables and the ‘car package’ was used to calculate the F-statistics as well as the p-values. In the data for the variable sugar content in adults, one outlier was removed. This variable in adults was boxcox transformed to meet the assumptions of normality of the residuals and homogeneity of variances.

### Appendix S3: Results of the bioenergetic response variables

In larvae, DNP exposure did not affect the fat content whereas low food decreased the fat content (~ -33 %) (Table S3.1., Fig S3.1.A). Sugar content was not affected by DNP exposure but was decreased by low food (~ -26 %) (Table S3.1., Fig S3.1.B). While DNP exposure did not affect the protein content, protein content was reduced by the low food level (Table S3.1., Fig S3.1.C).

In adults, fat content as well as sugar content were not affected by either the DNP or the food level. DNP exposure decreased the protein content slightly (Table S3.2., Fig S3.2.C) while low food had no effect.

Table S3.1. Results of LMs testing for the effects of DNP exposure and food level on the bioenergetic variables of the larvae of the damselfly *Lestes viridis.*

| Effect | Fat content | | | Sugar content | | | Protein content | | |
|---|---|---|---|---|---|---|---|---|---|
| | $df_1,df_2$ | F | p | $df_1,df_2$ | F | p | $df_1,df_2$ | F | p |
| DNP | 1,41 | 2.07 | 0.16 | 1,41 | 3.02 | 0.090 | 1,41 | 0.0008 | 0.978 |
| Food level | 1,41 | 18.49 | 0.00010 | 1,41 | 10.96 | 0.0019 | 1,41 | 16.55 | 0.00021 |
| DNP x Food level | 1,41 | 2.81 | 0.10 | 1,41 | 1.33 | 0.26 | 1,41 | 0.010 | 0.92 |

Table S3.2. Results of LMs testing for the effects of DNP exposure and food level on the bioenergetic variables of the adults of the damselfly *Lestes viridis.*

| Effect | Fat content | | | Sugar content | | | Protein content | | |
|---|---|---|---|---|---|---|---|---|---|
| | $df_1,df_2$ | F | p | $df_1,df_2$ | F | p | $df_1,df_2$ | F | p |
| DNP | 1,25 | 0.045 | 0.83 | 1,24 | 0.034 | 0.85 | 1,25 | 4.30 | 0.049 |
| Food level | 1,25 | 0.0074 | 0.93 | 1,24 | 0.36 | 0.55 | 1,25 | 1.70 | 0.20 |
| DNP x Food level | 1,25 | 0.58 | 0.45 | 1,24 | 0.32 | 0.58 | 1,25 | 1.73 | 0.20 |

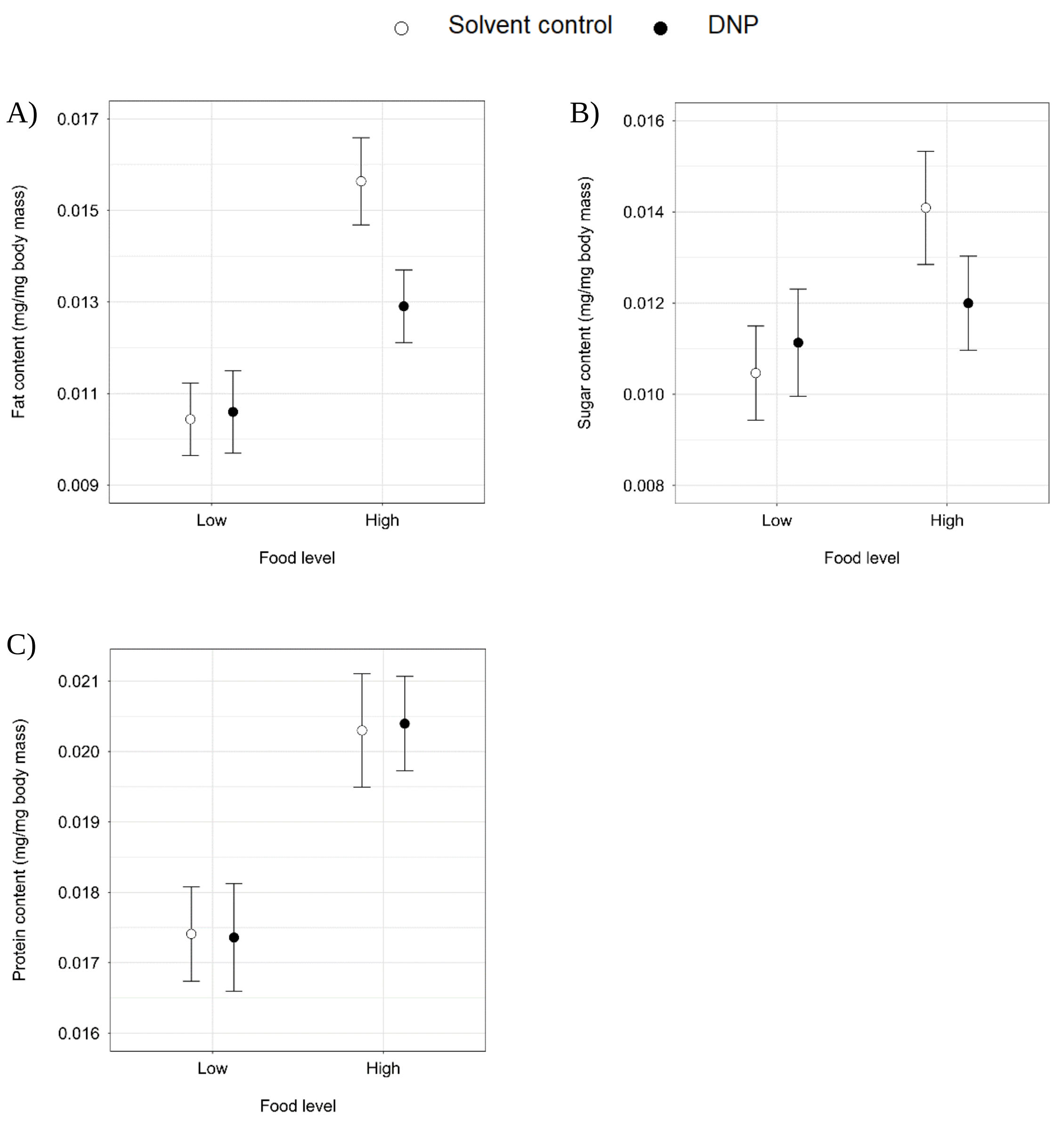


Figure S.3.1. The effect of DNP exposure and food level on the bioenergetic variables of the larvae damselfly *Lestes viridis*: (A) Fat content, (B) Sugar content, (C) Protein content. The means ± 1 SE are given.

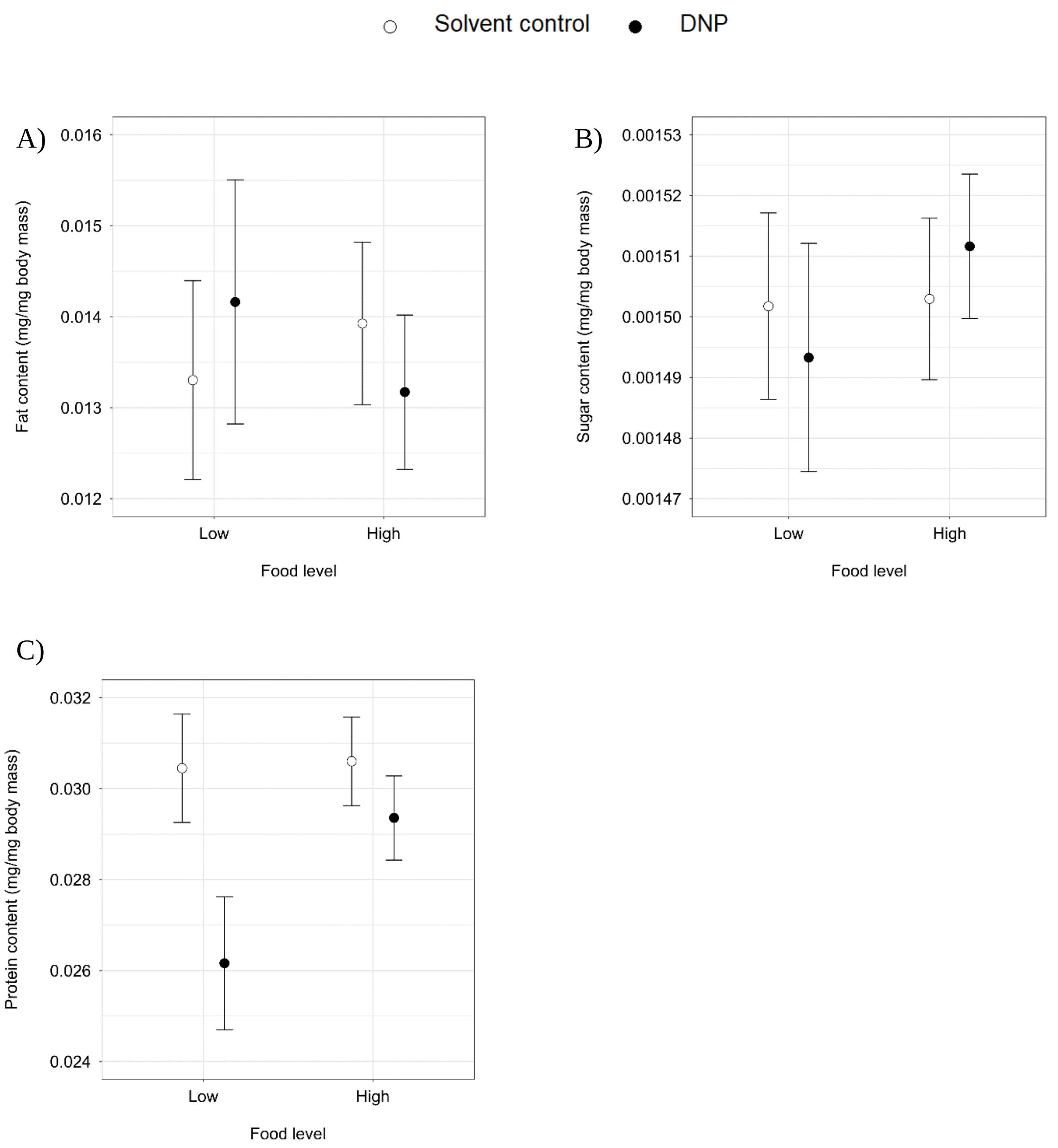


Figure S3.2. The effect of DNP exposure and food level on the bioenergetic variables of the adult damselfly *Lestes viridis*: (A) Fat content, (B) Sugar content, (C) Protein content. The means ± 1 SE are given

### Appendix S4: The full statistical tables and figures of the behavioural variables

Table S4.1. Results of GLMs testing for the effects of DNP exposure and food level on behavioural variables of the larvae of the damselfly *Lestes viridis*.

| Effect | Larval feeding strikes | | | Larval head orientations | | | Larval walks | | |
|---|---|---|---|---|---|---|---|---|---|
| | df | $\chi^2$ | p | df | $\chi^2$ | P | df | $\chi^2$ | p |
| DNP | 1 | 17.00 | <0.0001 | 1 | 11.33 | 0.00076 | 1 | 13.25 | 0.00027 |
| Food level | 1 | 10.31 | 0.0013 | 1 | 49.01 | <0.0001 | 1 | 40.65 | <0.0001 |
| DNP x Food level | 1 | 34.75 | <0.0001 | 1 | 27.85 | <0.0001 | 1 | 28.35 | <0.0001 |

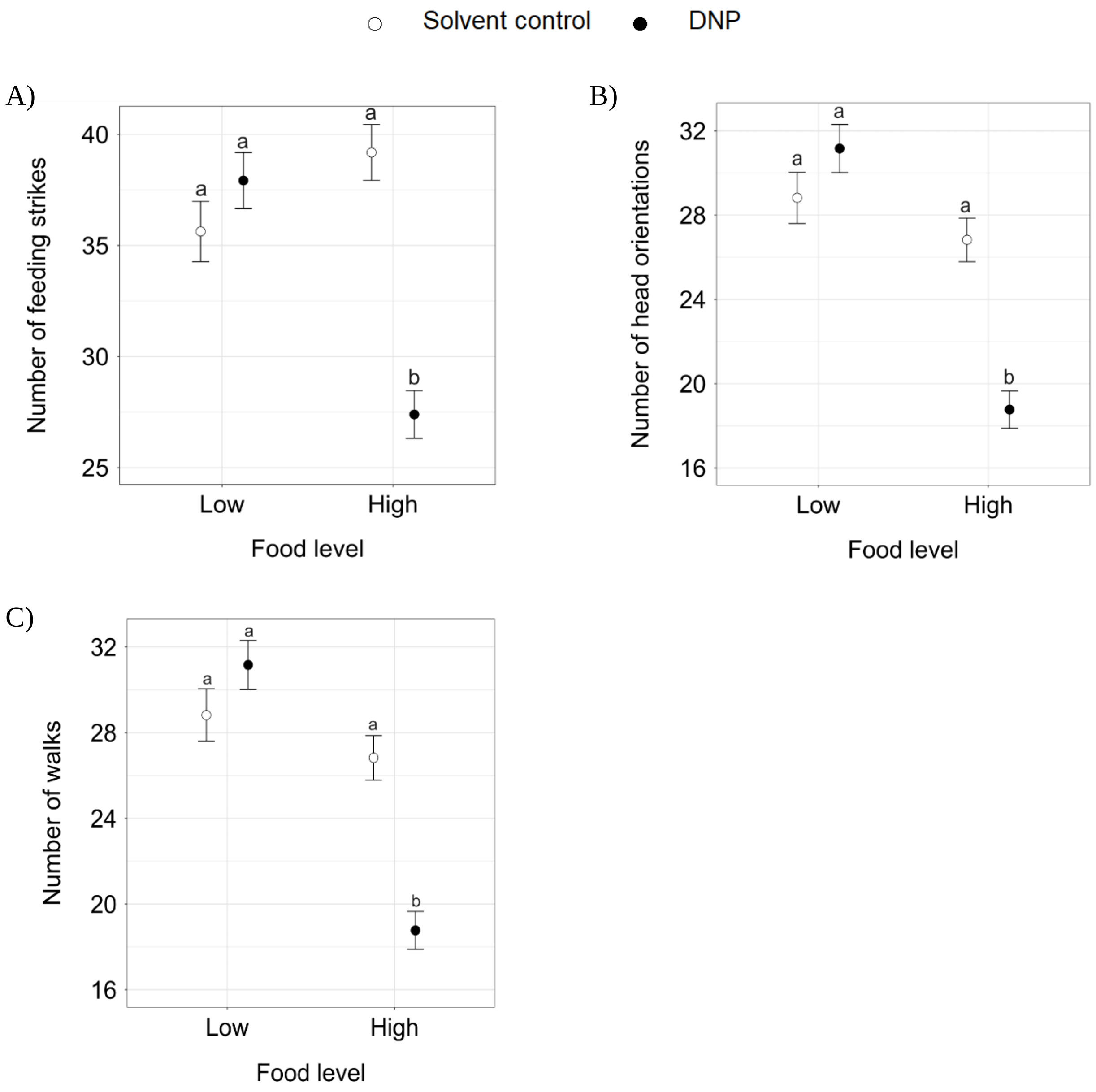


Figure S4.1. Effects of DNP exposure and food level on the behaviour of larvae of the damselfly *Lestes viridis*: numbers of (A) feeding strikes, (B) head orientations and (C) walks. All behavioural variables were corrected for larval mass. The means ± 1 SE are given. Different letters denote significant differences based on pairwise contrasts from the significant DNP × food level interaction.

**Appendix S5: The full statistical tables and figures of the physiological variables**

Table S5.1. Results of LMs testing for the effects of DNP exposure and food level on physiological variables of the larvae of the damselfly *Lestes viridis*.

| Effect | Energy available (Ea) | | | Energy consumed (Ec) | | | Cellular Energy Allocation (CEA) | | |
|---|---|---|---|---|---|---|---|---|---|
| | $df_1,df_2$ | F | p | $df_1,df_2$ | F | p | $df_1,df_2$ | F | p |
| DNP | 1,41 | 1.00 | 0.32 | 1,41 | 0.60 | 0.44 | 1,41 | 1.31 | 0.26 |
| Food level | 1,41 | 18.02 | 0.0001 | 1,41 | 5.13 | 0.03 | 1,41 | 2.41 | 0.13 |
| DNP x Food level | 1,41 | 1.73 | 0.20 | 1,41 | 0.41 | 0.52 | 1,41 | 0.65 | 0.43 |
| | MDA (Oxidative damage) | | | ATP/ADP | | | | | |
| | $df_1,df_2$ | F | p | $df_1,df_2$ | F | p | | | |
| DNP | 1,41 | 0.33 | 0.57 | 1,75 | 0.47 | 0.50 | | | |
| Food level | 1,41 | 9.20 | 0.0042 | 1,75 | 0.49 | 0.48 | | | |
| DNP x Food level | 1,41 | 2.74 | 0.11 | 1,75 | 0.50 | 0.48 | | | |

Table S5.2. Results of LMs testing for the effects of DNP exposure and food level on physiological variables of the adults of the damselfly *Lestes viridis*.

| Effect | Energy available (Ea) | | | Energy consumed (Ec) | | | Cellular Energy Allocation (CEA) | | |
|---|---|---|---|---|---|---|---|---|---|
| | $df_1,df_2$ | F | p | $df_1,df_2$ | F | p | $df_1,df_2$ | F | p |
| DNP | 1,25 | 2.48 | 0.13 | 1,25 | 4.53 | 0.043 | 1,25 | 0.93 | 0.35 |
| Food level | 1,25 | 1.12 | 0.30 | 1,25 | 8.44 | 0.0076 | 1,25 | 4.58 | 0.042 |
| DNP x Food level | 1,25 | 0.05 | 0.83 | 1,25 | 2.77 | 0.11 | 1,25 | 1.80 | 0.19 |

| Effect | MDA (Oxidative damage) | | | ATP/ADP | | |
|---|---|---|---|---|---|---|
| | $df_1,df_2$ | F | p | $df_1,df_2$ | F | p |
| DNP | 1,25 | 0.0001 | 0.99 | 1,57 | 3.65 | 0.061 |
| Food level | 1,25 | 0.055 | 0.82 | 1,57 | 3.72 | 0.059 |
| DNP x Food level | 1,25 | 0.0017 | 0.97 | 1,57 | 0.46 | 0.50 |

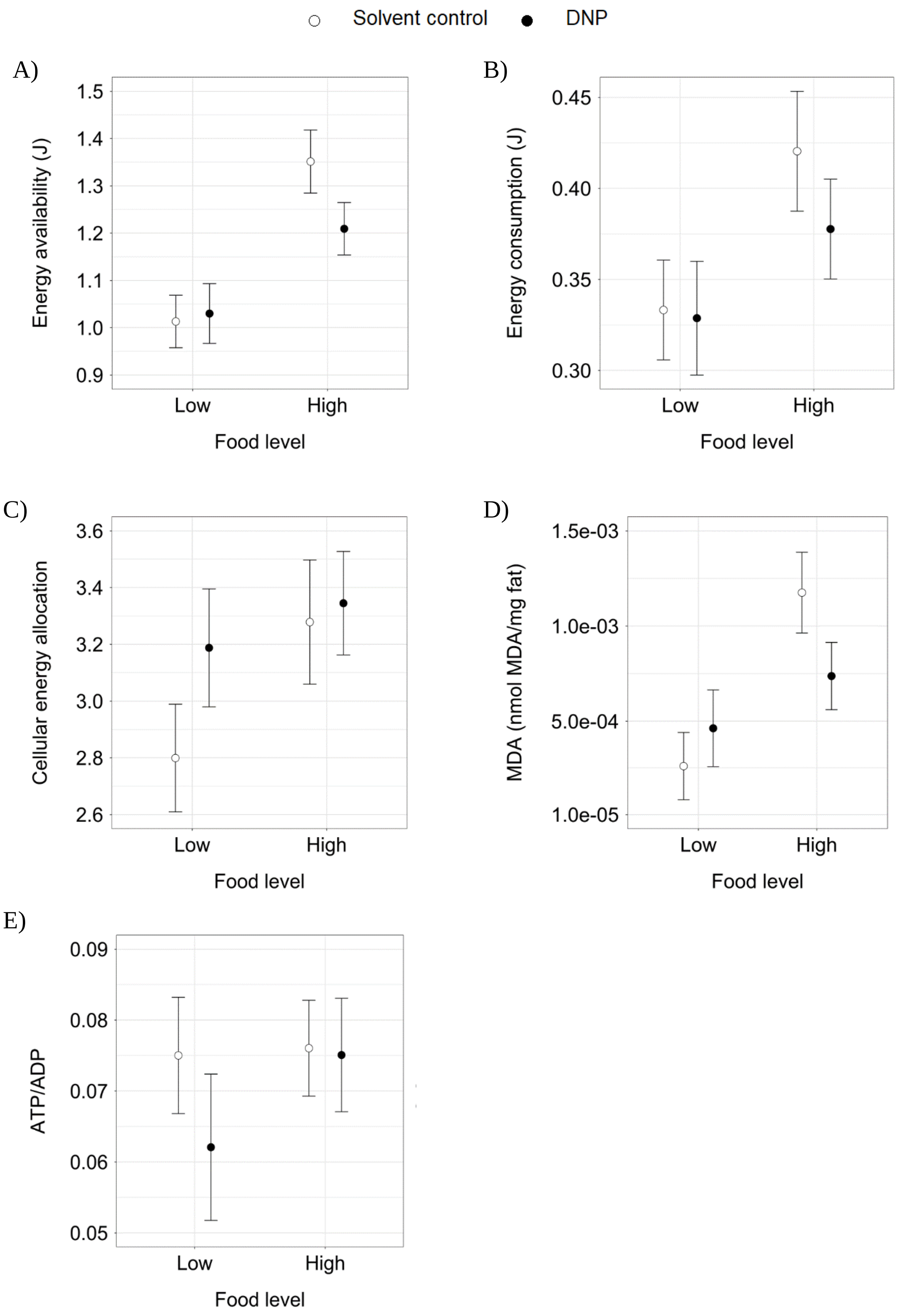
Solvent control
DNP
A)
Energy availability (J)
1.5
1.4
1.3
1.2
1.1
1.0
0.9
Low
High
Food level
B)
Energy consumption (J)
0.45
0.40
0.35
0.30
Low
High
Food level
C)
Cellular energy allocation
3.6
3.4
3.2
3.0
2.8
2.6
Low
High
Food level
D)
MDA (nmol MDA/mg fat)
1.5e-03
1.0e-03
5.0e-04
1.0e-05
Low
High
Food level
E)
ATP/ADP
0.09
0.08
0.07
0.06
0.05
Low
High
Food level

Figure S5.1. Effects of DNP exposure and food level on the physiological variables of the larvae of the damselfly *Lestes viridis*: (A) available energy (Ea), (B) consumed energy (Ec), (C) cellular energy allocation (CEA), (D) malondialdehyde (MDA) levels and (E) the ATP/ADP ratio. The means ± 1 SE are given.

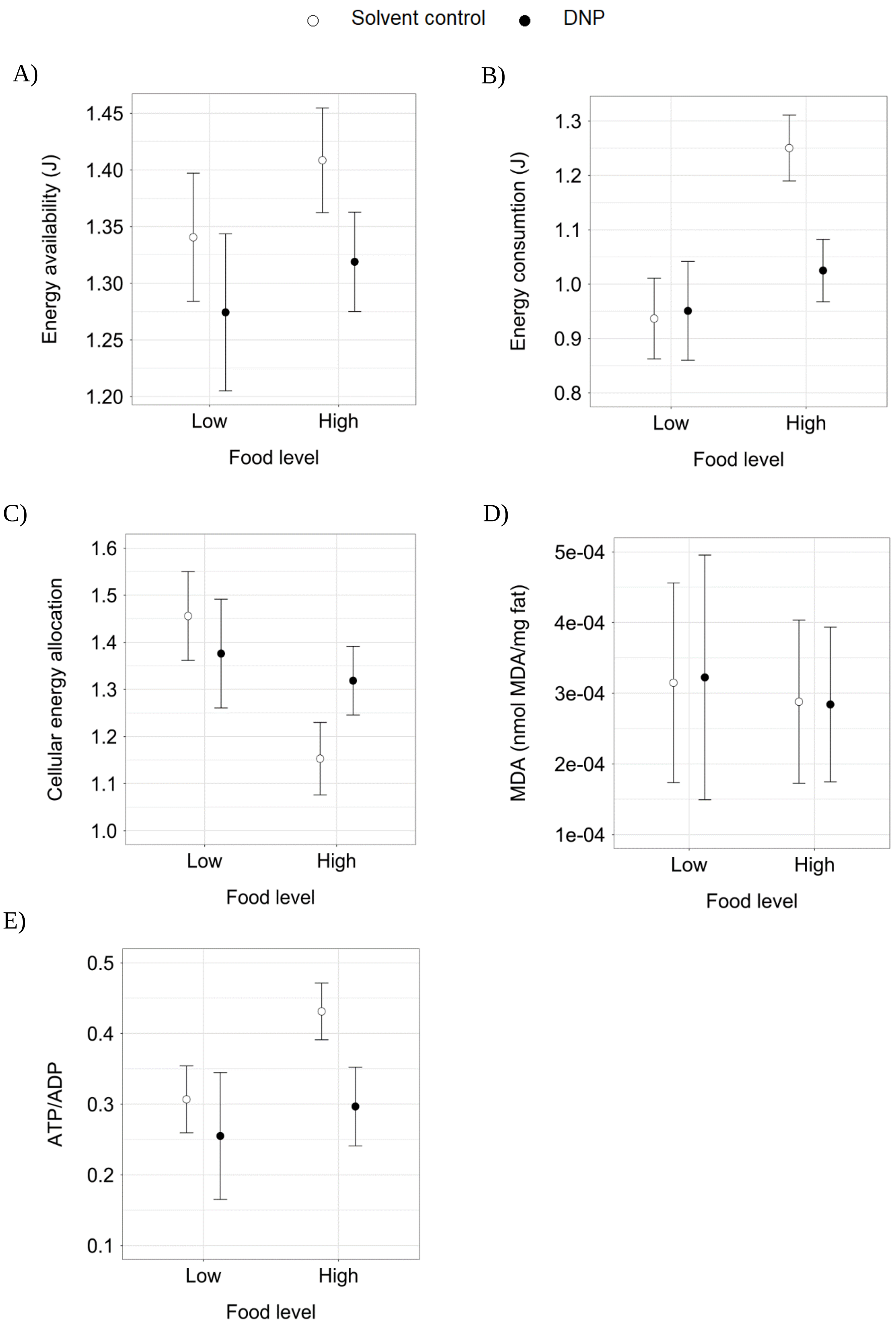

Solvent control
DNP
A)
Energy availability (J)
1.45
1.40
1.35
1.30
1.25
1.20
Low
High
Food level
B)
Energy consumtion (J)
1.3
1.2
1.1
1.0
0.9
0.8
Low
High
Food level
C)
Cellular energy allocation
1.6
1.5
1.4
1.3
1.2
1.1
1.0
Low
High
Food level
D)
MDA (nmol MDA/mg fat)
5e-04
4e-04
3e-04
2e-04
1e-04
Low
High
Food level
E)
ATP/ADP
0.5
0.4
0.3
0.2
0.1
Low
High
Food level

Figure S5.2. Effects of DNP exposure and food level on the physiological variables of adults of the damselfly *Lestes viridis*: (A) available energy (Ea), (B) consumed energy (Ec), (C) cellular energy allocation (CEA), (D) malondialdehyde (MDA) levels and (E) the ATP/ADP ratio. The means ± 1 SE are given.

**Appendix S6: Application of the independent action model for the survival-related variables**

An independent action (IA) model was used to determine whether the interaction between stressors was additive, antagonistic or synergistic for the survival-related variables (as suggested by Schäfer & Piggott, 2018). If the combined effect of the stressors is smaller than the effect predicted by the null model, the interaction is antagonistic (Côté et al., 2016; Schäfer & Piggott, 2018). Synergistic interactions are detected when the combined stressor effect is larger than the effect predicted by the null model while an additive interaction has no larger or smaller effect (Côté et al., 2016; Schäfer & Piggott, 2018).  To apply the IA model, we followed the protocol of Coors & de Meester (2008) for the variables larval survival, metamorphosis success and adult lifespan.

First, the proportional effects of the stressors were determined by using the following equation: $E_i = \frac{(e_i - e_{control})}{(e_{max} - e_{control})}$. In this formula, the $e_i$ represents the single-stressor effect and $e_{control}$ the effect of the (stress-free) control which are both expressed in absolute units. The $e_{max}$ represents the maximum possible single stressor effect, which is for larval survival and metamorphosis success zero and for adult lifespan 1. Second, we calculated the predicted joint effects using the equation $E_{joint} = 1 - \prod^{i} * (1 - E_i)$. Third, we compared the predicted joint effects with the observed joint effects. Therefore, we transformed $E_{joint}$ back to absolute units ($e_{joint} = E_{joint} * (e_{max} - e_{control}) - e_{control}$). Fourth, we constructed a 95 % confidence interval for the observed effect, expressed in absolute units, to assess if the observed effect and the predicted joint effect are significantly different from each other. An additive interaction between stressors is concluded if the predicted effect is in the 95 % confidence interval of the observed effect, while a synergistic interaction is concluded if the predicted effect is larger than the upper limit of the confidence interval (Coors & de Meester, 2008).

Furthermore, we also determined the strength of the interaction effect by calculating the model deviation ratio (MDR) whereby the predicted effect is divided by the observed effect (following Shahid et al., 2019). An MDR value of one represents an additive interaction effect while a higher MDR value represents a synergistic interaction effect. The higher the value, the stronger the synergism. Vice versa, when the MDR value is smaller than one, the interaction effect is antagonistic. The lower the value, the stronger the antagonism.

The IA model identified a synergistic interaction between the stressors DNP and low food for the three survival-related variables: larval survival, metamorphosis success and adult lifespan (Table S5.1.). Indeed, the MDR value for all the variables was higher than one (Table S5.1.). Notably, the synergism was the strongest for adult lifespan (MDR value of 12.80 vs other MDR values < 2.5).

Table S6.1. Results of the IA model for determining the interaction type between the stressors DNP and low food. This model was used for the three survival-related variables: larval survival, metamorphosis success and adult lifespan.

| | Predicted | Observed | 95 % CI observed | Interaction effect | Model deviation ratio |
|---|---|---|---|---|---|
| Larval survival | | | | | |
| DNP x Low food | 0.54 | 0.29 | [0.24, 0.34] | Synergistic | 1.86 |
| Metamorphosis success | | | | | |
| DNP x Low food | 1.02 | 0.45 | [0.29, 0.61] | Synergistic | 2.27 |
| Adult lifespan | | | | | |
| DNP x Low food | 12.80 | 1 | [0.95, 1.02] | Synergistic | 12.80 |
